# Endocytic clathrin-coated vesicles are targeted for selective autophagy during osmotic stress

**DOI:** 10.1101/2025.09.16.676479

**Authors:** Jonathan Michael Dragwidge, Matthieu Buridan, Julia Kraus, Thibault Kosuth, Clément Chambaud, Lysiane Brocard, Klaas Yperman, Evelien Mylle, Michaël Vandorpe, Dominique Eeckhout, Geert De Jaeger, Roman Pleskot, Amélie Bernard, Daniël Van Damme

## Abstract

Plants frequently encounter hyperosmotic stress due to drought and salinity, leading to rapid water loss, reduced turgor pressure, and decreased cell volume. This contraction drastically alters plasma membrane tension, a physical parameter that must be strictly maintained to support mechanosignaling and cell expansion. However, the mechanisms by which plants adjust their membrane surface area to match a shrinking cell volume remain poorly understood. Here, we identify selective autophagy dependent degradation of plasma membrane-derived clathrin-coated vesicles (CCVs) in response to hyperosmotic shock. This pathway involves the recruitment of the endocytic TPLATE complex (TPC) to autophagosomes in an osmotic-stress dependent manner. Through correlative light and electron microscopy (CLEM) and electron tomography (ET), we provide ultrastructural evidence of the physical association of CCVs with autophagosome membranes. These autophagosomes contain endocytic machinery, including TPC and clathrin, and are targeted to the vacuole. Mechanistically, we show that the conserved ATG8-interacting motifs (AIMs) in the AtEH1/Pan1 and AtEH2/Pan1 TPC subunits interact with ATG8, suggesting that they facilitate the recruitment of CCVs to autophagosomes. Using time-lapse imaging, we demonstrate that the acute induction of autophagy is precisely coupled to the reduction in cell volume under hyperosmolar conditions. Our results suggest that endocytic removal of excess plasma membrane to maintain membrane tension and cellular integrity is coupled to TPC-mediated CCV-phagy. These findings reveal a homeostatic mechanism that enables plants to adapt to the challenges of drought and salinity.

**Significance Statement:** All living cells must maintain the physical integrity of their outer membrane. How cells can adapt their surface area to respond to rapid changes in cell volume, such as those caused by drought or salt stress, remains a fundamental question in biology. We identify a mechanism in plants involving autophagy – a cellular recycling pathway – where plasma membrane-derived vesicles are targeted for degradation following salt or osmotic stress. This pathway involves the TPLATE complex, an evolutionary ancient and essential endocytic complex in plants, which directly interacts with the autophagy protein ATG8. This discovery reveals how plants adapt to osmotic stress by modulating their membrane properties, supporting a framework for future improvement of crop resilience in response to salinity and drought.

## Introduction

Macroautophagy (hereafter autophagy), is a conserved eukaryotic pathway, which degrades intracellular components to maintain cellular homeostasis and to recycle building blocks. Autophagy is activated by environmental and developmental cues to selectively remove proteins, aggregates, and damaged or superfluous organelles. In plants, autophagy is integral for development and for adaptation to diverse abiotic stresses, including nutrient deprivation, drought, salinity, and endoplasmic reticulum stress (1–4).

Autophagy proceeds through the *de novo* formation of double-membraned vesicles, termed autophagosomes, which sequester cargo and deliver it to the vacuole for degradation. This process is orchestrated by autophagy-related (ATG) proteins that coordinate autophagosome nucleation, membrane expansion, and fusion (5). ATG8 proteins are central to this pathway: upon lipidation they associate with the expanding autophagosomal membrane, promoting its growth while recruiting cargo. Selective cargo recognition depends on interactions between ATG8 and cargo receptors containing ATG8-interacting motifs (AIMs), which confer specificity by linking particular proteins or organelles to the autophagosome membrane (6). Several plant autophagy receptors have been characterised, including NBR1, which recognises aggregated ubiquitinated cargo (7, 8), TSPO, which promotes degradation of aquaporins (9), ATI1/2, which mediate plastid turnover (10, 11), and CESAR/ERC which function in heat stress recovery (12, 13). However, the repertoire of receptors and cargo that mediate stress adaptation remains incompletely defined.

The TPLATE complex (TPC) is an evolutionary ancient adaptor complex that is essential for clathrin-mediated endocytosis in plants (14, 15). As an octameric assembly, TPC contributes to membrane deformation and cargo recognition (16, 17), and cooperates with clathrin, adaptors including adaptor protein complex 2 (AP-2), and membrane phospholipids to generate clathrin-coated vesicles (18). The TPC subunits AtEH1/Pan1 and AtEH2/Pan1 are implicated in lipid binding (16, 19), cargo interaction (19), and phase separation (20), and were previously linked to autophagy during prolonged nutrient starvation (21). Whether TPC participates more broadly in autophagy pathways, particularly under acute stress conditions, remains unknown.

Hyperosmotic shock imposes a physical challenge to the plasma membrane. Rapid water efflux reduces cell volume (22), lowering membrane tension (23, 24), and creates a sudden excess of plasma membrane surface area (25). To restore homeostasis, this excess membrane must be removed via internalization. Because membrane tension is a primary determinant of cell expansion and mechanosignaling, its regulation is likely under tight homeostatic control (26). Endocytosis rates are modulated by osmotic stress (27), yet how the excess plasma membrane material is ultimately processed remains unclear. Given that salt and osmotic stress also trigger robust autophagic activity (28, 29), we questioned whether these two pathways intersect to manage the sudden influx of endocytic material.

Here, we identified an autophagy pathway in which TPC-labelled clathrin-coated vesicles are selectively recruited to autophagosomes during hyperosmotic stress. Using live-cell imaging, correlative light and electron microscopy, and quantitative proteomics in *Arabidopsis thaliana*, we demonstrate that the EH-domain subunits of TPC contain functional AIM motifs that directly interact with ATG8. These results are consistent with TPC having a role as a selective autophagy receptor to target clathrin-coated vesicles for vacuolar degradation. This mechanism facilitates the rapid turnover of endocytic vesicles, plasma membrane, and associated cargo, and is tightly coupled to stress-induced changes in cell volume and membrane tension. Our findings reveal that plants employ CCV-phagy as a rapid homeostatic rheostat to restore plasma membrane integrity under osmotic challenge.

## Results

### TPC is recruited to autophagosomes following salt and osmotic stress

The endocytic TPC has been implicated in an autophagy pathway at ER-PM contact sites during long-term nutrient starvation (21), yet its function in other autophagy inducing stresses is unclear. To investigate this, we examined whether the outer TPC subunit AtEH1/Pan1, and the ‘inner core’ subunit TPLATE associate with the autophagosome marker ATG8 using native promoter complemented *A. thaliana* reporter lines. We tested commonly used autophagy stresses, including Target of Rapamycin (TOR) inhibition, salt stress, and osmotic stress. We combined these stresses with co-treatment with the actin depolymeriser Latrunculin B (LatB) to stabilise autophagosomes in the cytosol by slowing their fusion with the vacuole. We found that both AtEH1/Pan1 and TPLATE puncta associated with ATG8 labelled phagophores following NaCl and sorbitol treatment (Fig. 1a-d). In contrast, AZD8055 treatment did not significantly induce association of TPC with ATG8 compared to LatB alone (Fig. 1a-d). These data suggest that the association of TPC with ATG8 is triggered by specific environmental stimuli linked to salt and/or osmotic stress.

**Fig. 1.**
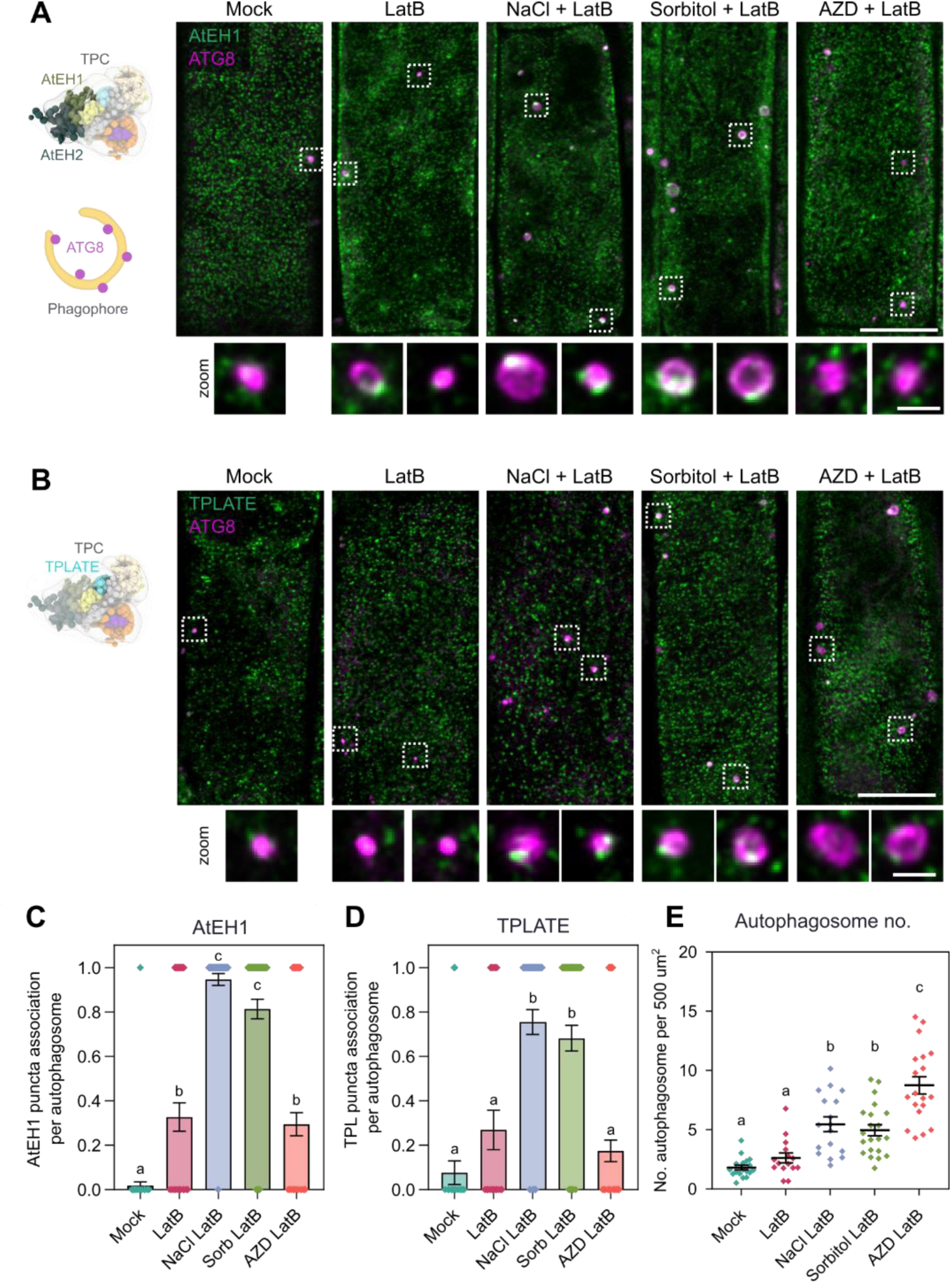
TPC associates with autophagosomes during salt and osmotic stress. (**A-B**) TPC model (16) positioning the TPC outer subunits AtEH1 and AtEH2, and the inner core subunit TPLATE. Airyscan images show association of AtEH1-GFP and TPLATE-GFP puncta with autophagosomes (mCherry-ATG8e) in *A. thaliana* root epidermal cells. Seedlings were imaged 20 minutes after treatment with the actin depolymeriser Latrunculin B, salt (NaCl), sugar (sorbitol), or the TOR kinase inhibitor AZD8055. Insets show high-contrast zoomed images of AtEH and TPLATE puncta associated with ATG8 labelled autophagosomes. (**C-D**) Quantification of association of AtEH1 and TPLATE puncta on autophagosomes. Bars indicate mean ± SEM; n = 23–80 autophagosomes. **(E)** Quantification of average autophagosome number per root (normalised by area) from epidermal cells. Bars indicate mean ± SEM; n = 15–19 roots. **(C – E)** Statistical significance was determined by one-way ANOVA with Tukey’s HSD post hoc test (α = 0.05); different letters indicate significant differences between samples. Scale bars: 10 µm, 1 µm (inset).

To test whether ionic (e.g. Na^+^) or osmotic components were responsible for the observed effects, we examined the association of AtEH1/Pan1 with ATG8 using other ionic compounds (Fig. S1a). AtEH1/Pan1 association with ATG8 was similar regardless of the ionic species (Fig. S1b), indicating that the association was due to osmotic, not ionic effects. Additionally, NaCl and sorbitol treatment induced similar amounts of autophagosomes (Fig. 1e), further supporting a primary role that osmotic stress triggers autophagosome formation, and that this process is distinct from autophagy induction following TOR inhibition. As hyperosmolarity has been linked to increases in free Ca^2+^ (30), we examined whether free Ca^2+^ influences autophagosome formation. Ca^2+^ channel inhibition using Lanthanum chloride (LaCl_3_) did not alter AtEH1/Pan1-ATG8 association (Fig. S1c-d8). We therefore conclude that our observations are not a consequence of altered cytosolic Ca^2+^ levels. Lastly, we performed time-lapse imaging of seedlings following NaCl treatment, and we identified AtEH1/Pan1 puncta in stable association with a phagophore/autophagosome throughout autophagosome maturation (Fig. S2). Together these data imply that both outer and inner core TPC subunits are selectively recruited to autophagosomes in an osmotic stress dependent manner.

### CCVs physically associate with autophagosomes upon osmotic stress

To further characterize the nature of TPC association with autophagosomes following sorbitol treatment, we performed ultrastructural studies using correlative light and electron microscopy (CLEM). Using CLEM, we identified ATG8-labelled phagophores and autophagosomes (Fig. 2a-b), although we could not detect AtEH1/Pan1-GFP fluorescence, likely due to low signal intensities following fixation. Electron tomography (ET) reconstructions of autophagic structures revealed distinct clathrin-coated vesicles (CCVs), which were physically connected to the phagophore/autophagosome membrane (Fig. 2b and S3a). Quantification of the frequency of CCV association to autophagosomes between sorbitol and AZD treated roots (31) showed that CCV-autophagosome association is much more prevalent following sorbitol treatment compared to nutrient starvation mimicking conditions (Fig. 2c). These data are consistent with our live cell imaging data (Fig. 1), and imply that CCV association with autophagosomes is not a general feature of autophagy, but is a stress specific feature. We also observed enlarged autophagosome-multi-vesicular body (MVB) hybrid compartments (i.e. amphisomes) with associated CCVs (Fig. S3a-b). These structures are consistent with previous reports that autophagosomes can fuse with MVBs before their subsequent delivery to the vacuole (32, 33). Collectively, these findings indicate that endocytic-derived, TPC-labelled CCVs associate with autophagosomes specifically upon osmotic stress.

**Fig. 2.**
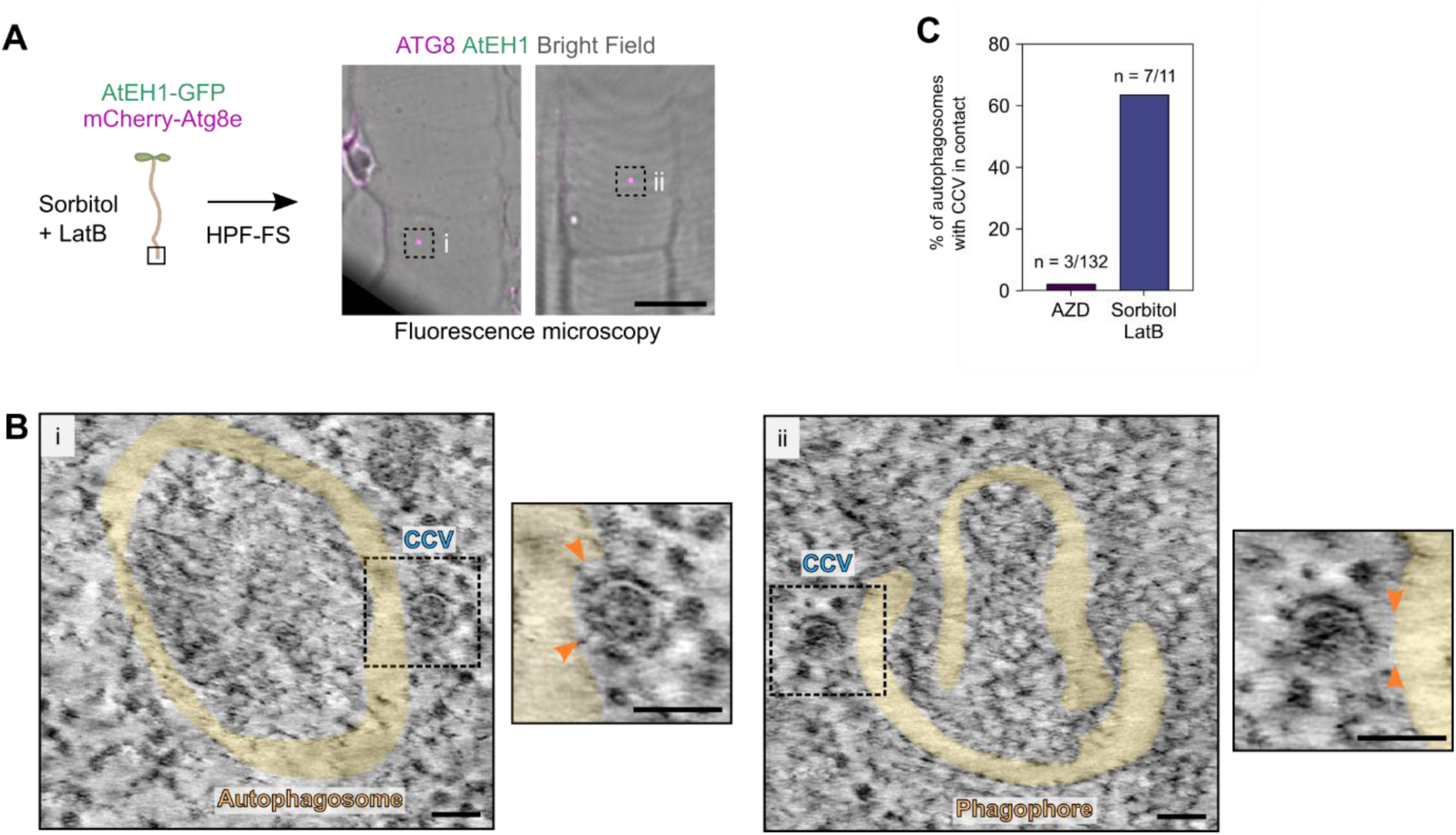
CLEM-ET reveals that CCVs are physically associated to autophagosomes. **(A)** Correlative light and electron microscopy (CLEM) from high-pressure frozen (HPF), freeze substituted (FS) *A. thaliana* roots co-expressing AtEH1-GFP and mCherry-ATG8e. The image shows punctate ATG8 signal in root epidermal cells taken from a single 150 nm ultrathin section. **(B)** Electron tomography reconstructed slices from the identified phagophore/autophagosomes via CLEM in (A). Autophagosome membranes (false coloured in yellow) are shown. Clathrin-coated vesicles that are physically attached to the phagophore are indicated and shown enlarged (insets, arrowheads). (**C**) Proportion of electron tomography reconstructed autophagosomes with clathrin-coated vesicles attached from plants treated with Sorbitol + LatB (this study) or AZD alone (31). Scale bars: 10 µm (a), 50 nm (b).

### Plasma membrane cargo and endocytic machinery are delivered to the vacuole during osmotic stress

If endocytic CCVs are degraded by autophagy, then the endocytic machinery and the internalised plasma membrane cargo should be associated with autophagosomes. To test this, we performed proteomic analysis of immuno-precipitated autophagic compartments enriched from sorbitol treated *A. thaliana* seedlings (Fig. 3a). We identified plasma membrane receptors and associated proteins enriched in the ATG8 bound fraction (Fig. 3b), implying that CCVs containing plasma membrane proteins associate with autophagosomes. Furthermore, core endocytic machinery including subunits of the adaptor protein 2 (AP-2) complex, and the scission protein dynamin-related protein 2b (DRP2b) were significantly enriched in the ATG8 fraction. To confirm that CCVs with their associated cargo and accessory proteins- are delivered to the vacuole via autophagy, we visualised vacuolar autophagic bodies by inhibiting vacuolar acidification using Concanamycin A (ConcA) (34). Upon sorbitol and ConcA co-treatment we observed prominent and progressive accumulation of vacuolar bodies containing clathrin (clathrin-light chain), TPC (AtEH1/Pan1 and TASH3), and dynamin (DRP1a) (Fig. 3c and 3e). These data indicate that endocytic machinery and clathrin are degraded in the vacuole. As a control, we examined AP-1 and AP-4, which respectively label a TGN secretory, and vacuolar trafficking pathway (35) (Fig. 3d). AP-1 showed minimal accumulation in vacuolar bodies, while AP-4 accumulated in the vacuole (Fig. 3c-d), likely via MVB-autophagosome intermediate compartments (32, 33). These findings confirm that endocytic-derived CCVs are delivered to the vacuole for degradation following osmotic stress.

**Fig. 3.**
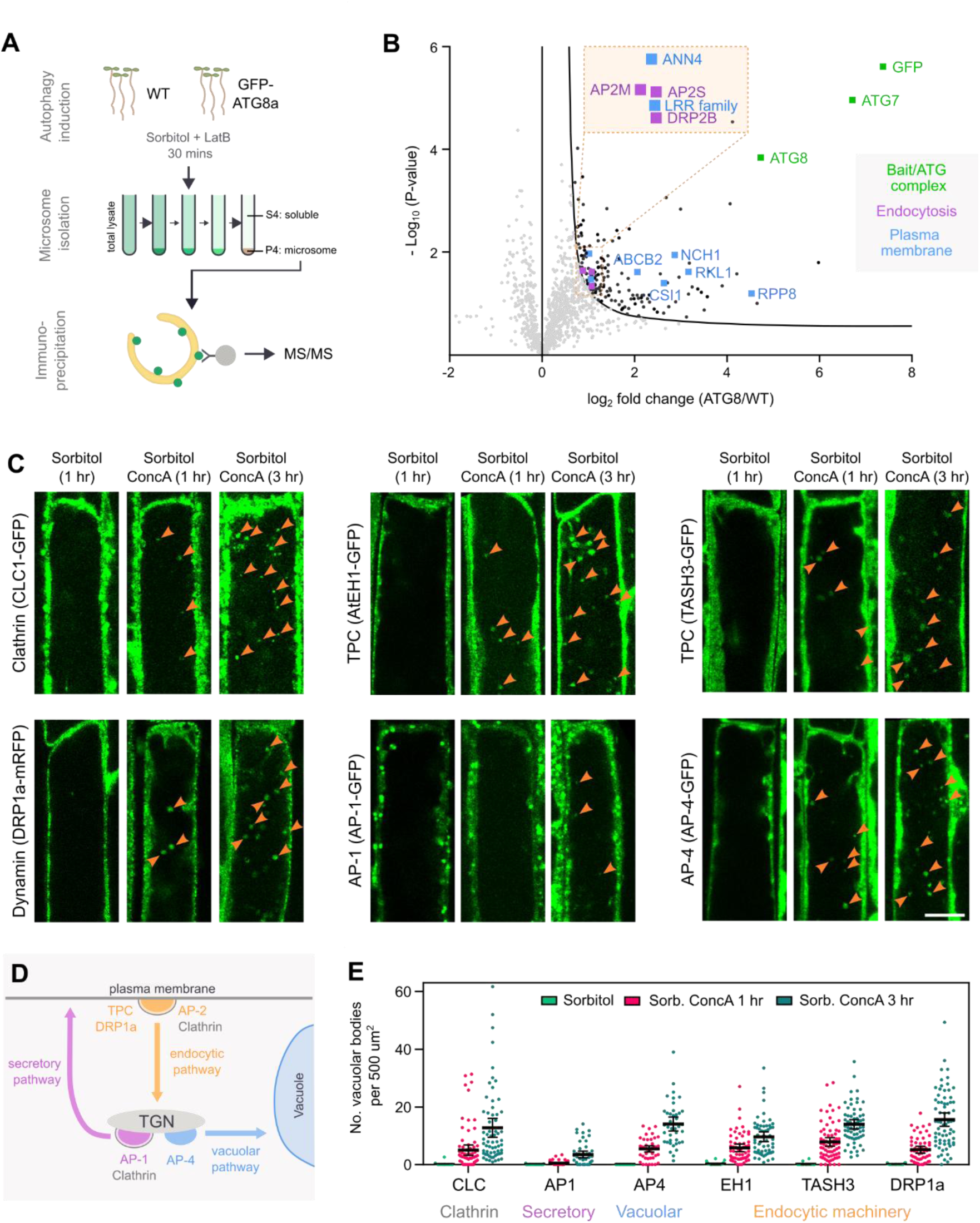
Plasma membrane cargo and endocytic machinery are delivered to the vacuole following osmotic stress induced autophagy. **(A)** Workflow for microsome isolation, autophagosome pull-down, and proteomics. **(B)** Volcano plot showing proteins enriched in the GFP-ATG8a fraction compared to WT. Significant enrichment of plasma membrane (blue), endocytic (purple), and bait (green) proteins are highlighted. **(C)** Confocal images of root epidermal cells from the indicated trafficking-related reporters following mock (sorbitol) treatment, or co-treatment with sorbitol and Concanamycin A (ConcA). Vacuolar autophagic bodies are visible following ConcA treatment (arrowheads). Images are contrast adjusted to visualise vacuolar signal. **(D)** Simplified schematic of major trafficking pathways. **(E)** Quantification of vacuolar bodies. Bars indicate mean ± 95% CI; n = 26–38 (sorbitol) or 50–73 (ConcA) root cells. Scale bars: 10 µm.

### AtEH1/Pan1 and AtEH2/Pan1 directly interact with ATG8

Selective autophagy involves the specific recognition of cargo for degradation, canonically via the presence of ATG8 interacting motifs (AIMs), which bind the hydrophobic pocket of ATG8, thereby mediating cargo recruitment to autophagosomes (1). We previously reported that AtEH1/Pan1 contains an acidic region, which comprises a putative AIM motif consisting of [W/F/Y]–X–X–[L/I/V] (21) (Fig. 4a). Evolutionary analysis revealed that most AtEH/Pan1 homologs contain two tandem AIMs, except for dicotyledon EH1/Pan1 proteins (e.g. AtEH1/Pan1), which lack the second AIM (Fig. 4b and S4). To test whether the putative motif that is present in the C-terminal intrinsically disordered region 3 (IDR3) of AtEH1/Pan1 is a functional AIM, we performed a phase separation partitioning assay in *N. benthamiana*. ATG8 partitioning into AtEH1/Pan1 droplets was dependent on the presence of IDR3, which contains the AIM motif (Fig. 4c). Furthermore, Alphafold3 modelling predicts that putative AIMs in AtEH1/Pan1 and AtEH2/Pan1 interact with the hydrophobic pocket of ATG8 (Fig. 4d-g). To directly test the AIM-ATG8 interaction, we performed a pull-down experiment using recombinantly purified ATG8 and biotinylated peptides containing the AIMs and mutated AIMs of AtEH1/Pan1 and AtEH2/Pan1 (Fig. 4h). As a positive control, we used AIM peptides of the known ATG8-interactor ATG1 (36). All AIM-containing peptides directly interacted with ATG8, similarly to the ATG1 AIM peptide, while AIM mutated variants failed to do so (Fig. 4i). Collectively, these experiments confirm that AtEH/Pan1 proteins physically interact with ATG8 via their AIMs. These data imply that the AtEH/Pan1 TPC subunits could function as selective autophagy receptors, linking endocytic CCVs to autophagosomes.

**Fig. 4.**
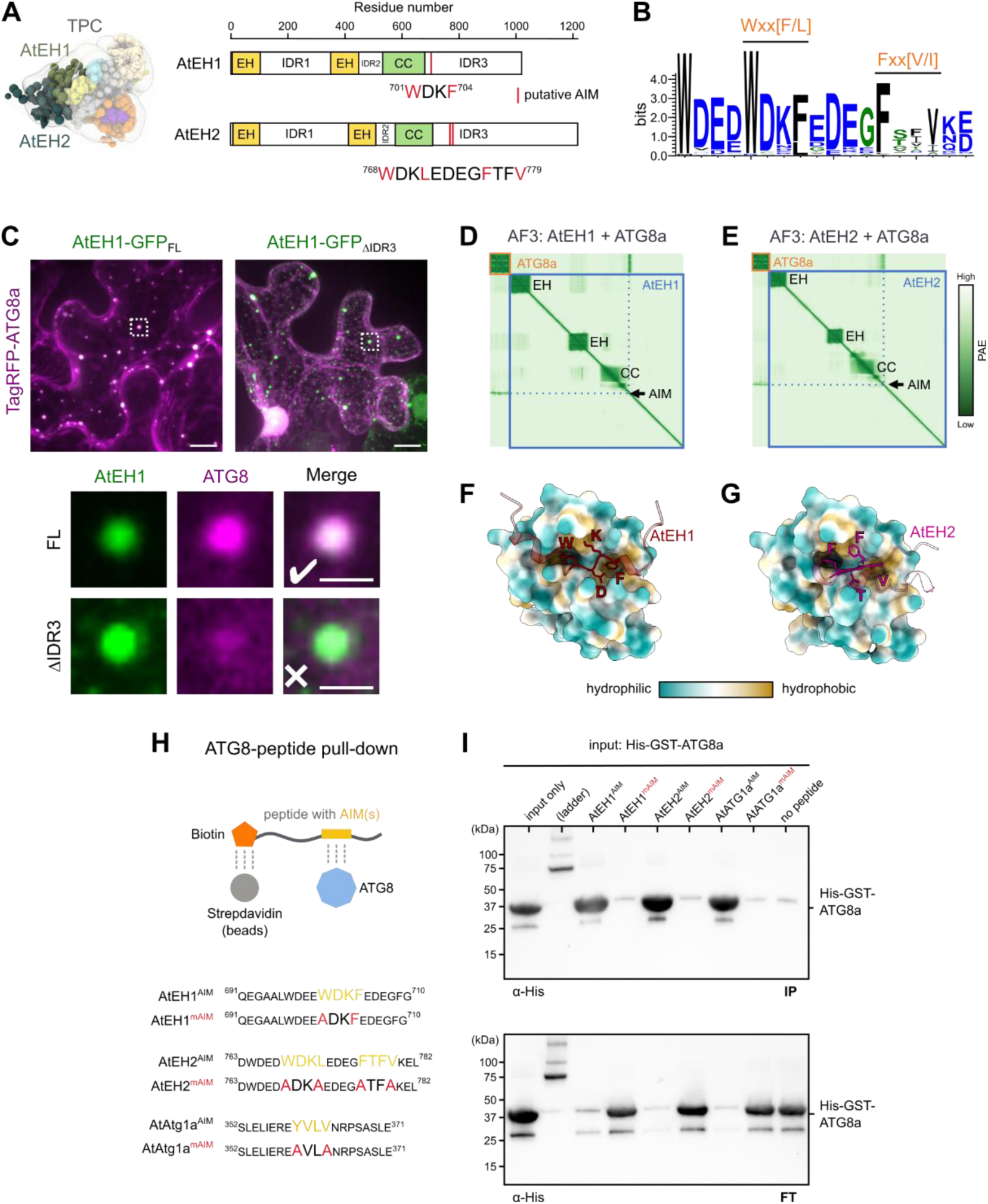
Both AtEH/Pan1 proteins physically interact with ATG8 via classical AIMs. **(A)** Model of AtEH1/Pan1 and AtEH2/Pan1 within the TPC (16), and schematic representation of the AtEH proteins with indication of the position of the putative ATG8 interaction motifs (AIMs). **(B)** Logo plot of the AIM region from 61 plant EH proteins across evolution (see Fig. S4). (**C**) Partitioning recruitment assay of AtEH1-GFP and TagRFP-ATG8a in *N. benthamiana* epidermal cells. The inset shows presence or lack of recruitment of ATG8 to AtEH1-GFP condensates. (**D-G**) Alphafold3 prediction of the interaction of ATG8 and AtEH1/AtEH2. **(D-E)** Predicted alignment error (PAE) plots of ATG8a and AtEH1/AtEH2 showing high confident interaction in the AIM. (**F-G**) Model of AtEH1(WxxF) and AtEH2 (FxxV) AIMs binding to the LDS hydrophobic pocket of ATG8a. (**H**) Schematic representation of the ATG8-peptide^AIM^ pull-down experiment, with the sequence of peptides and AIMs indicated. AtATG1a was used as a positive control. (**I**) Western blot showing the immunoprecipitate (IP) and flow-through (FT) of samples following ATG8-peptide pull-down. ATG8 is present in the IP fraction in peptides containing the AIMs, while remains in the FT in peptides with mutated AIMS (mAIM). Scale bars: 10 µm, 2 µm (inset).

### Autophagy and cell volume changes are inversely correlated during osmotic stress

Hyperosmotic stress triggers a reduction in cell volume, which is coupled to a corresponding decrease in plasma membrane tension (23, 24) (Fig. 5a), impairing membrane function. We therefore asked whether CCV-dependent autophagic degradation of the plasma membrane could function to support turn-over of excessive plasma membrane and thereby assist in restoring membrane tension following a decrease in cell volume. To test this, we performed live-cell imaging of *A. thaliana* roots expressing a plasma membrane marker and an autophagy marker following hyperosmotic shock using a custom-made microfluidics setup on a vertical confocal microscope (Fig. 5b). We obtained cell volume measurements of individual cells via segmentation, which we combined with autophagosome quantification to obtain high-resolution dynamics during hyperosmotic shock and recovery. Hyperosmotic shock produced an immediate drop in cell volume (to ∼ 90% of pre-shock), which gradually recovered over 72 minutes (Fig. 5c-e, S6, Video S1). In contrast, autophagosomes were induced as soon as 4 minutes post treatment, peaked around 16 minutes, and returned to baseline levels by approximately 64 minutes. Thus, autophagosome formation and cell volume are inversely correlated. We also performed longer-term imaging of seedlings after transfer to hyper-osmotic agar media, which produces an initial hyperosmotic shock and prolonged hyperosmotic stress as the seedlings grow on the agar (Fig. S6, Video S2). High levels of autophagy were triggered by hyperosmotic shock, which also continued at lower levels for several hours after transfer, likely representing local sensing of osmotic stress. The strong inverse correlation between cell volume and autophagy, combined with the above data strongly suggest that autophagy actively contributes to restore membrane tension following osmotic stress by degrading endocytosed CCVs.

**Fig. 5.**
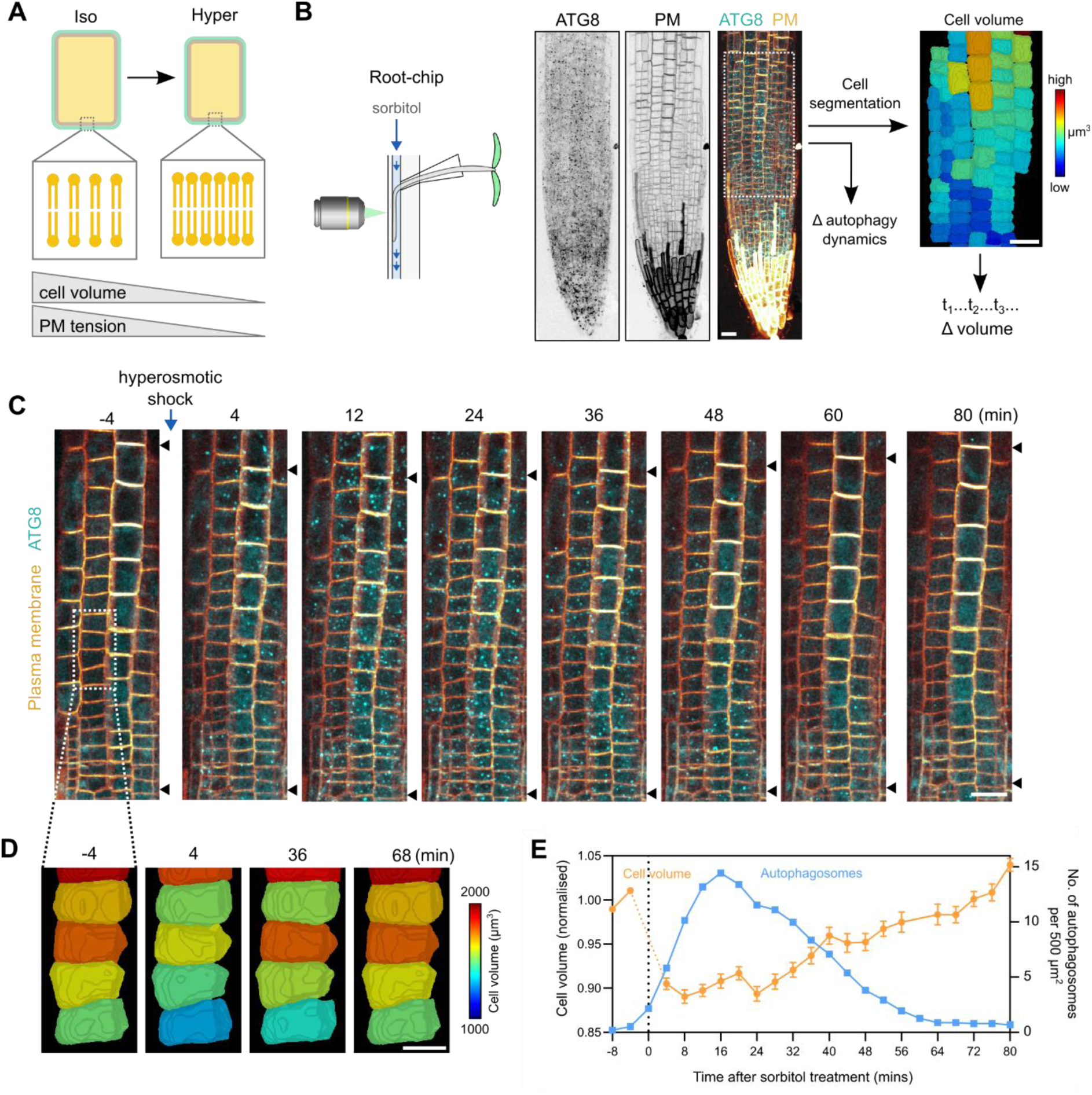
Autophagy and cell volume changes are inversely correlated during hyperosmotic shock recovery. **(A)** Schematic indicating the coupling of cell volume and plasma membrane tension at iso- and hyper-osmotic conditions. **(B)** Schematic of the imaging setup. *A. thaliana* seedlings were mounted on a root-chip microfluidics chamber and pump connected to a vertical confocal microscope. Confocal images show *A. thaliana* root cells with co-labelling of autophagosomes (YFP-ATG8a), and plasma membrane (mCherry-SYP122) which enables cell volume segmentation. **(C)** Time-lapse images before and after hyperosmotic shock. Arrowheads indicate region the region used for analysis, which undergoes compaction following hyperosmotic shock. **(D)** Zoomed in segmented volume reconstruction showing cell volume recovery over time from the indicated cells (dashed lines). **(E)** Quantification of autophagosome number and cell volume during the time-course. Cell volume data is mean ± SEM; n = 31 cells. Scale bars: 20 µm (C), 10 µm (D).

## Discussion

In this study, we have uncovered a function of TPC in autophagy-mediated degradation of clathrin-coated vesicles following hyperosmotic stress. We identified that hyperosmotic stress induced autophagosomes contain TPC and other endocytic machinery, which are actively delivered to- and degraded in the vacuole. Thus, we propose that this mechanism effectively removes plasma membrane lipids and cargo following hyperosmotic stress. Through high-spatiotemporal imaging of hyperosmotic shock, we found that autophagy is induced immediately after cell volume loss and the corresponding decrease in membrane tension. We therefore propose that autophagic degradation of CCVs functions to maintain plasma membrane homeostasis by removing excess membrane to restore membrane tension after hyperosmotic induced cell shrinkage.

This pathway likely acts in parallel with enhanced endocytic flux following osmotic stress (27, 37). However, while standard endocytosis primarily redistributes lipids and integral membrane proteins within the endomembrane system (38), autophagic degradation of plasma membrane-derived CCVs results in the effective removal of membrane material, as illustrated by our observation that plasma membrane proteins accumulate in vacuolar bodies. This is a notable advantage in the context of long-term adaption to drought and salt stress conditions, which reduce water availability and cause cell shrinkage. This model is supported by our observation that autophagy is activated by both acute and prolonged hyperosmotic stress. Consequently, our study provides a potential cellular mechanism to explain the long-observed induction of autophagy by drought and salt (28, 29, 32). Beyond volume regulation, it remains to be determined whether membrane tension-mediated pathways involving mechanotransduction (39, 40), or polarity-driven growth (41) are influenced by TPC-mediated CCV-phagy.

Our work also expands the role of TPC and the AtEH/Pan1 subunits in endomembrane trafficking. AtEH subunits have diverse functions ranging from cargo recognition (19), to membrane-dependent biomolecular condensation (20), and are linked to the actin cytoskeleton, ER-PM contact sites, and autophagy (21). Our findings extend this view by showing that AtEH/Pan1 subunits contain effective ATG8 binding AIM motifs, consistent with a role as selective receptors to degrade clathrin-coated vesicles via autophagy following hyperosmotic stress. The conservation of AtEH/Pan1 AIMs across diverse plant species implies that this mechanism is evolutionarily ancient and has an important function in plants.

How AtEH/Pan1 proteins function as selectively autophagy receptors to recognise CCVs for autophagic degradation remains unclear. The cell must differentiate between CCVs that need to be delivered to the TGN/EE via canonical endocytosis, and those destined for autophagy. Our findings indicate that endocytic CCVs that are degraded by autophagy retain cytosolic endocytic proteins, including multiple TPC subunits, dynamin, and clathrin. This is unusual, as canonical endocytic vesicles undergo uncoating after vesicle scission from the plasma membrane, resulting in the removal of clathrin and accessory proteins (including TPC and dynamin) (42–44), which enables vesicle fusion with the TGN/EE. This finding could indicate that the retention of TPC on CCVs and presence of the AtEH/Pan1 AIMs could act as an ‘eat me’ signal to trigger CCV-phagy, while CCVs where the TPC is removed following scission follow the canonical pathway and fuse with the TGN/EE. Osmotic stress mediated increase in cell crowding is known to trigger conformational changes in IDRs (45, 46), and may facilitate exposure of the AIMs in the AtEH1/AtEH2 IDR to enable recognition by ATG8.

Our proteomic analysis did not detect strong enrichment of TPC in ATG8 pull-downs, suggesting that TPC may only loosely associate with CCVs on autophagosomes. Nonetheless, both clathrin and TPC subunits have been previously observed in the vacuole following nutrient starvation and ConcA treatment (21), and our experiments similarly detected clathrin, TPC, and endocytic cargo accumulation in the vacuole after hyperosmotic stress. These findings confirm that endocytic vesicles can be delivered and degraded in the vacuole. It is unclear whether vesicles targeted for autophagy carry specific cargo that is actively degraded for adaption against osmotic stress. Since TPC subunits including AtEH1/Pan1 can recognise plasma membrane cargo (17, 19), future work could define whether specific cargo are preferentially degraded and how they mediate stress adaption.

### Limitations of the study

While our results demonstrate a strong correlation between cell volume and autophagy induction after hyperosmotic shock, we lack direct membrane tension measurements. Consequently, while at short timescales membrane tension changes are coupled to cell volume, the precise kinetics of membrane tension during hyperosmotic shock and recovery remain to be defined. Future studies using tension-sensitive FRET sensors will be required to resolve this.

Furthermore, while our data identify functional AIM motifs in both AtEH/Pan1 subunits that physically interact with ATG8, the specific requirement of these motifs for CCV-phagy during osmotic stress remains to be genetically established. Likely functional redundancy between AtEH1/Pan1 and AtEH2/Pan1 at the level of AIM sequences, combined with the male sterility phenotypes of individual *eh1* and *eh2* mutants (14, 21), presents a significant challenge for genetic complementation studies. Definitive confirmation of their roles as autophagy receptors will require the generation of viable double-complemented lines expressing AIM-mutated variants of both proteins.

## Materials and methods

### Plant material and growth conditions

*Arabidopsis thaliana* accession Columbia-0 (Col-0) plants were used for all experiments. Seeds were surface sterilised by chlorine gas and grown on ½ strength Murashige and Skoog (½ MS) medium containing 0.8% (w/v) agar, pH 5.8, with 1% sucrose, or without sucrose for microfluidics experiments. Seedlings were stratified for 48 h at 4°C in the dark, and transferred to continuous light conditions at 21 °C in a growth chamber. Imaging was performed on 5-6-day old seedlings unless otherwise indicated. The *Arabidopsis* lines expressing *pCLC1:CLC1-GFP* (47), *p35S:DRP1a-mRFP* (48), *pAP1M:AP1M2-GFP* (49)*, pAP4M*:*AP4M-GFP* (50)*, pCLC1:CLC1-GFP*, *pH3.3:TASH3-GFP* (17), *pUBQ10:GFP-ATG8a* (51), and *pLAT52:TPLATE-GFP × p35S:mCherry-ATG8E* (21) were described previously. Double-marker lines expressing *pAtEH1:AtEH1-mGFP* (20) and *p35S:mCherry-ATG8e* (52); and *pUBQ10:3XmCherry-SYP122* (53) and *pUBQ10:YFP-ATG8a* (54), were generated by crossing the respective single-reporter homozygous lines.

### Transient expression in N. benthamiana

*N. benthamiana* plants were grown under long-day conditions (16h light, 8 hr dark, 100 μmol PAR, 21 °C) in soil supplemented with Saniflor Osmocote Pro. Leaves were co-infiltrated with Agrobacterium tumefaciens (strain GV3101) (OD_600_ = 0.5) containing plasmids UBQ10:AtEH1_FL_-mGFP or UBQ10:AtEH1_1-684_-mGFP (20), and pUBQ:3xHA-TagRFP-AtAtg8a, together with the p19 silencing inhibitor. At 3–4 days post-infiltration (dpi), epidermal cells were imaged using a PerkinElmer UltraView spinning-disk confocal system mounted on a Nikon Ti inverted microscope. Images were captured with an ImagEM CCD camera (Hamamatsu C9100-13) and Volocity software using a 60× NA 1.2 water-immersion objective. Sequential acquisition was performed using a 488-nm laser with a 500–550 nm band-pass filter for GFP, and a 561-nm laser with a 570–625 nm band-pass filter for RFP.

### Chemical and salt treatments

For chemical treatments seedlings were incubated in 6-well plates containing ½ MS media supplemented with Latrunculin B (4 μM) or Concanamycin A (1 μM) from 1000× DMSO stocks. For autophagy induction, seedlings were treated with ½ MS solutions containing sorbitol (200 mM), NaCl (100 mM) for the indicated durations. These concentrations were chosen as they stimulated autophagy and cell volume shrinkage without compromising cellular integrity of inducing plasmolysis.

### Confocal microscopy and image analysis

Confocal imaging was performed on a Zeiss LSM880 Airyscan microscope using a 63× Oil (NA 1.4) Apo, or 40× Oil (NA 1.3) Apo objective. Airyscan acquisition was performed in RS mode with GFP [BP 495-550] and RFP [LP570] emission filters, with subsequent processing in Zeiss ZEN Blue software. Timelapse imaging of AtEH1 and ATG8 was performed on a Leica DMI8 spinning-disk microscope equipped with a Yokogawa CSU-W1 head, 491-nm and 561-nm diode lasers, and a Photometrics Prime 95B camera, and a 63× oil objective.

To quantify TPC-ATG8 association, ROIs of autophagosomes were defined at their medial plane from Z-stacks (4 μm range, 1 μm spacing) using ImageJ. AtEH1/TPLATE puncta were scored as associated if they were on, or touching the autophagosomal membrane. Autophagosome density was determined using the LoG detector in the TrackMate plugin (ImageJ), and total counts per ROI were normalised to the root surface area (μm^2^).

For quantification of autophagic bodies following ConcA treatment, time-lapse images were acquired in confocal mode using a GaAsP detector. Autophagic bodies were identified by their characteristic rapid stochastic motion within the vacuole, distinguishing them from other cytosolic compartments. Autophagic bodies were detected in Trackmate from ROIs of the vacuole from individual cells, and normalised to the surface area (um^2^).

### Microfluidics and vertical microscope imaging

Microfluidic experiments were performed using custom chips and holders as previously described (55). Chip inlets were connected to a Fluigent system (Fluigent, Jena, Germany) with LineUp pressure pumps (Flow EZ™ 1000), flow sensors (Flow Rate Sensor M), and 15 ml reservoirs containing filtered 1/2 MS medium with or without 200 mM Sorbitol. TRITC-dextran 20 kDa (Sigma-Aldrich, 73766) was diluted 1:1000 in the treatment medium to monitor medium switching, which was achieved by setting the desired flow to 3 µL/min while the other inlet was set to 0 µL/min. Flow rates were controlled via the Oxygen software (Fluigent).

Seedlings (4–5 days old) were mounted in the chips and holders as previously described (55). Seedlings were placed on a custom vertical-stage Zeiss LSM 900 KAT confocal microscope before imaging and recovered for 20 min under a constant flow of 3 µl/min. Roots were imaged with a Zeiss Plan-Apochromat 20×/0.8 dry objective. GFP and mCherry were acquired sequentially with 488 nm and 561 nm lasers and GaAsP-PMT detectors (detection windows: 490–546 nm and 560–700 nm). Z-stacks (1 µm step size) were collected at 240-s intervals over 30 time points using a two-tile scan. Images were recorded at 1024 × 1024 pixels, 8-bit depth, with a pixel dwell time of 0.52 µs in bidirectional scanning mode. Acquisition settings were adjusted to prevent pixel saturation when projecting slices. External illumination was maintained throughout imaging. For long-term imaging the TipTracker script (56) was used to maintain the root tip in the field of view.

3D cell volume was quantified using the MorphoGraphX software package. Image stacks were first pre-processed in ImageJ (Fiji) for 3D drift correction. Corrected stacks were imported into MorphoGraphX and processed with ITK Gaussian smoothing. Segmentation was achieved via ITK watershed auto-seeding based on signal intensity. 3D meshes were generated using the Marching Cubes algorithm (cube size 0.5, smoothing 3). Individual cell volumes were extracted using the ‘Heat Map Classic’ tool.

### CLEM-ET

CLEM was performed as previously described (20, 57). Seedlings were incubated in ½ MS containing 200 mM sorbitol and 4 µM LatB for 20 minutes. Root tip segments were excised and cryofixed via high-pressure freezing (Leica EM ICE) using 20% BSA as a cryoprotectant. Freeze-substitution was conducted (Leica AFS2) in acetone containing 0.1% (w/v) uranyl acetate, followed by embedding in Lowicryl HM20 resin.

Ultra-thin sections (150 nm) were obtained using a Leica Ultracut microtome with a diamond knife and collected on copper mesh grids. Fluorescence was visualized on a Zeiss LSM 880 (40× oil-immersion objective, NA 1.3). Ultrastructural observations were performed using an FEI Tecnai Spirit (120 kV) equipped with an Eagle 4K×4K CCD camera or an FEI Talos F200S G2 (200 kV). ATG8-labeled autophagosomes were identified by correlating fluorescence signals with electron micrographs using cell boundaries as fiducial landmarks. Electron tomography tilt series were acquired from -65° to 65° at 1-degree increments, and reconstructed using the SIRT-like filter in the eTomo software in IMOD as previously described (20, 58).

### Protein expression and purification

HIS-GST-ATG8a protein was expressed in Escherichia coli BL21 (DE3) cells. Protein expression was induced by 0.4 mM isopropyl β-D-1-thiogalactopyranoside (IPTG) for 6 h at 30°C. Cells were collected by centrifugation and lysed with lysis buffer (50 mM Tris-HCl pH8, 500 mM NaCl, 5% glycerol, 50mM imidazole, 20mM glycine, Protease inhibitor (cOmplete ULTRA EDTA-free, Roche), 1/1000 benzonase). The bacteria were lysed by sonication, centrifuged (30 min, 40,000 x g), filtered through a Minisart NML GF filter (Sartorius), and loaded into a IMAC column (HiPrep IMAC FF 16/10). Proteins were eluted in reverse flow using a single step to 500mM imidazole, then loaded on a Hiload Superdex 200pg 16/600 equilibrated a SEC buffer (20mM HEPES 7.5, 150mM NaCl). Protein samples were eluted and stored in the SEC buffer at -70°C.

### ATG8-peptide pull-down

Biotinylated peptides of Biotin-Ahx-(Ahx)x4-Peptide (Genscript) were dissolved in DMSO at 5mg/mL, and diluted 5x in SEC buffer with 2mM DTT. Streptavidin sepharose high performance beads (Cytiva) were washed 3x with the SEC buffer. Peptides (15 nmol) were added to beads and incubated for 1h at 4°C, and then beads were washed 3x in the SEC buffer. HIS-GST-ATG8 (6 nmol) was added to beads and incubated for 1h at 4°C, and then beads were washed 4x in the buffer and the supernatant removed. Protein was eluted with Laemmli buffer by vortexing for 30 sec, and the beads were boiled at 75°C for 8 mins. Samples were loaded and separated on a 4–20% SDS–PAGE TGX gel (BioRad) and subsequently blotted on polyvinylidenedifluoride (PVDF; BioRad) using anti-HIS (Invitrogen, 1:1000). HRP-conjugated antibodies were detected using western lighting plus-ECL reagent (PerkinElmer).

### Autophagosome pull down

The autophagosome pull-down was performed as described (59), following microsome preparation as previously described (32). Briefly, approximately 10 mg of WT and GFP-ATG8a *Arabidopsis* seeds were grown on ½ MS plates vertically under continuous light for 6 d. Seedlings were transferred to 6 well plates containing 200 mM sorbitol and 4 uM LatB for 30 minutes, then ground at 4°C in GTEN-based buffer (10% glycerol, 30mM Tris pH 7.5, 150mMNaCl, 1 mM EDTA pH 8, 0.4M sorbitol, 5mM MgCl2, 1 mM Dithiothreitol (DTT), 1× liquid protease inhibitor cocktail (Sigma-Aldrich), and 1% Polyvinylpolypyrrolidon (PVPP)) in a 3:1 v/w ratio. Next, lysates underwent several centrifugation steps at 4 °C (10 min 1,000 g, 10 min 10,000 g, 10 min 15,000 g), where each time the supernatant was transferred. Protein concentration in S3 was normalised through BCA (Sigma-Aldrich) to ensure that an equal amount of protein was loaded before ultracentrifugation (60 min at 100,000 g at 4°C). The P4 microsome fraction was resuspended in GTEN buffer containing 0.1% Nonidet P-40, and samples were immunoprecipitated using GFP-Trap Magnetic Agarose beads (Chromotek). Following IP, samples were resuspended in 5% SDS in 50 mM TEAB pH 8.5 and incubated at 10 minutes at room temperature with mixing by pipetting to elute samples from beads.

### Proteomic analysis

Protein samples were analysed by 2-h data-independent acquisition (DIA) runs (60) on an Orbitrap Exploris 240 mass spectrometer. Raw data were processed with DIA-NN v2.0.2. A predicted spectral library was first generated from the *Arabidopsis thaliana* Araport11plus_DE2023.fasta database, against which the raw data were searched. This yielded 1,842 protein groups. Contaminants were removed, and groups were filtered to retain only protein groups supported by at least two unique peptides. The processed protein group file was imported into Perseus (61) for downstream statistical analysis. LFQ intensities were log2-transformed, and proteins were filtered to retain those with at least four valid values in one experimental group. Missing values were imputed from a normal distribution (width = 0.3, downshift = 2.0). Differential abundance analysis was performed using a two-sample *t*-test comparing ATG8-GFP pulldown versus wild-type control, with significance thresholds set at FDR = 0.05 and S0 = 0.5.

### Multiple sequence alignment and phylogenetic analysis

The identification and phylogenetic analysis of AtEH1 homologues used for AIM analysis is previously described (20).

### Structural modelling of the interaction between ATG8 and AtEH1/AtEH2

Alphafold3 (62) was used to predict interactions between ATG8 and AtEH1 or AtEH2 with full-length protein sequences as input. The top-scoring models were selected and visualised using UCSF ChimeraX (63).

### Statistics and reproducibility

Statistical analysis was performed using Graphpad Prism and Microsoft Excel. Significance criterion was set at a p value of <0.05. No statistical methods were used to predetermine sample size, but our sample sizes are similar to previously reported studies. Experiments were repeated at least twice, unless otherwise indicated. For comparisons between two groups, a two-tailed Student’s t-test was used. For multiple comparisons, statistical significance was determined by one-way or two-way ANOVA followed by Tukey’s honestly significant difference (HSD) post hoc test. The significance criterion was set at P<0.05.

### Data availability

All materials are available from the corresponding authors upon request.

## Acknowledgements

We thank Joop Vermeer, Ikuko Hara-Nishimura, and Tomoo Shimada for sharing published material. The authors would like to thank Iene Rutten (MeBioS KULeuven) and the VIB TechWatch FabLab project for the funding and expertise to develop the microfluidics setup on the vertical confocal microscope, Inge Verstraeten (UGent) and Matyas Fendrych (IEB Prague) for providing the Rootchips and for their expertise and advice in handling the microfluidics setup. We thank the VIB proteomics core facility for the Exploris 240 analysis. This work was supported by the Research Foundation – Flanders (FWO) research grant G017919N (D.V.D.), a FWO postdoctoral fellowship 12S7222N (J.M.D.) and a FWO long stay abroad grant V427024N (J.M.D.). This project also received funding from the European Research Council (ERC) under the European Union’s Horizon 2020 research and innovation program (grant agreement No 852136 to AB). Microscopy was done at the Bordeaux Imaging Center, a member of the national infrastructure France-BioImaging supported by the French National Research Agency (ANR-10-INBS-04).

## Author contributions

Conceptualisation: J.M.D., A.B., and D.V.D. Investigation: J.M.D., M.B., J.K., T.K., C.C., K.Y., E.M., M.V., D.E., and R.P. Writing: J.M.D., A.B., and D.V.D. Supervision: D.V.D., A.B., and G.D.J.

---

**Fig. S1.**
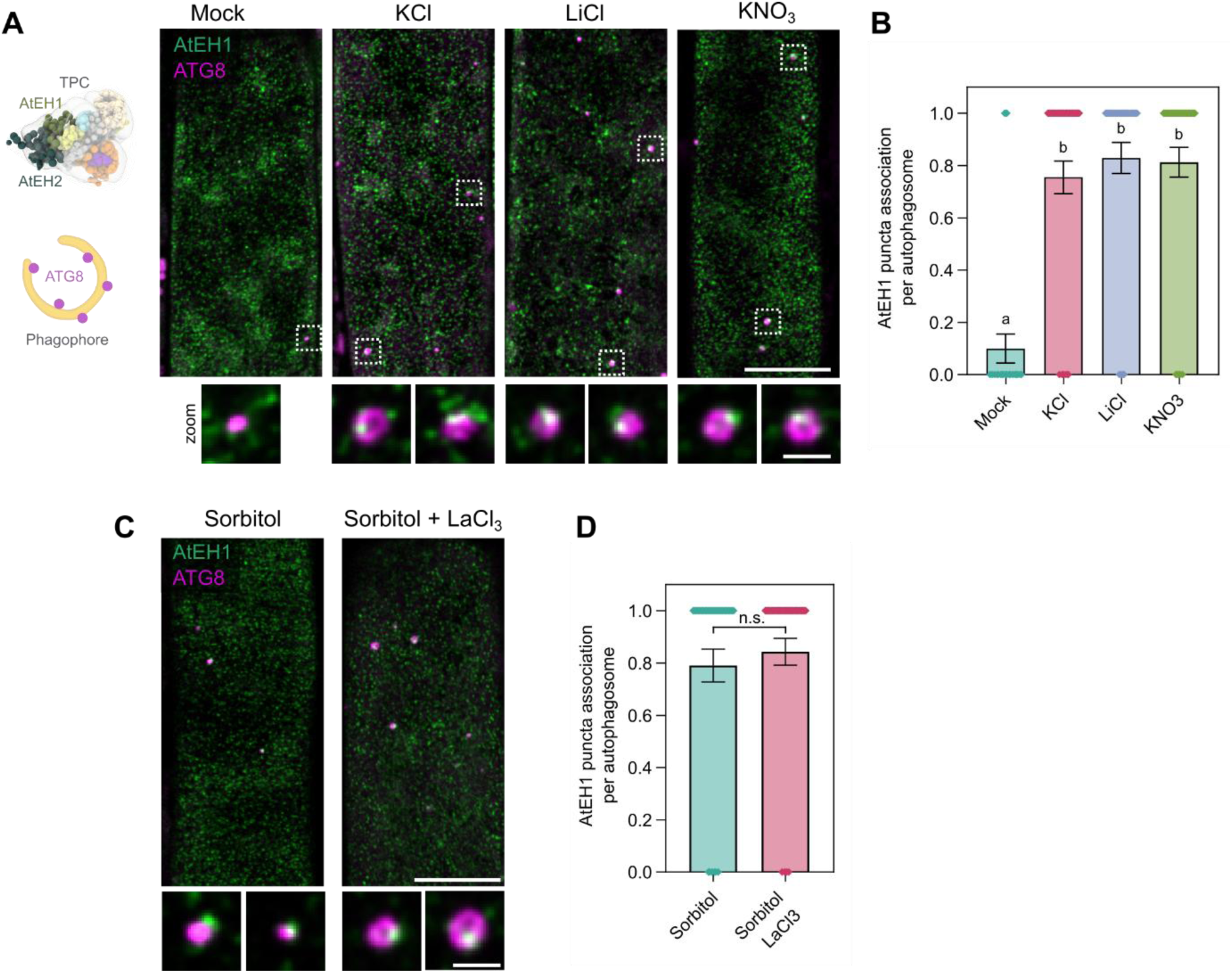
(Related to Fig. 1). AtEH1/Pan1 association with autophagosomes is not specific for NaCl, and is independent of plasma membrane Ca^2+^ channel activity. **(A)** Airyscan images of AtEH1-GFP and mCherry-ATG8e in *A. thaliana* root epidermal cells. Seedlings were imaged 20 minutes after treatment with the indicated salts (100 mM) together with Latrunculin B (LatB). **(B)** Quantification of association of AtEH1 puncta with autophagosomes. Bars indicate mean ± SEM; n = 30 – 49 autophagosomes. Statistical significance was determined by one-way ANOVA with Tukey’s HSD post hoc test (α = 0.05); different letters indicate significant differences between samples. **(C)** Airyscan images of AtEH1-GFP and mCherry-ATG8e in *A. thaliana* root epidermal cells after treatment with the Ca^2+^ channel blocker LaCl_3_. **(D)** Quantification of AtEH1 association with autophagosomes. Bars indicate mean ± SEM; n = 43 – 51 autophagosomes. p values: unpaired t-test with Welch’s correction; n.s. not significant. Scale bars: 10 µm, 1 µm (inset).

**Fig. S2.**
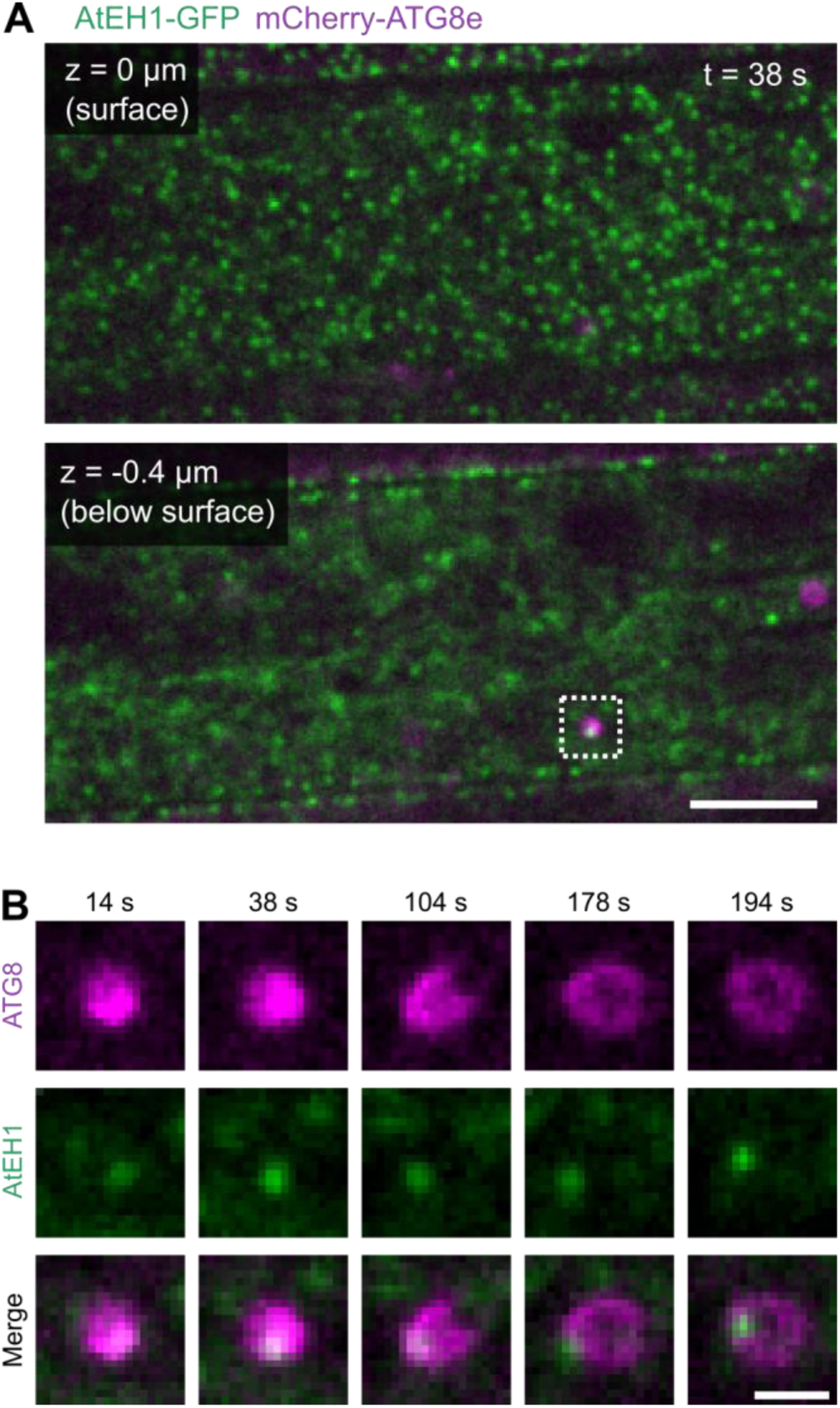
(Related to Fig. 1). Timelapse imaging of AtEH1/Pan1 and ATG8 during autophagosome progression. **(A-B)** Spinning-disk microscopy images of AtEH1-GFP and mCherry-ATG8e in *A. thaliana* root epidermal cells after 15 minutes treatment with 50 mM NaCl. **(A)** Single slices from the cell surface plane, and slightly under the cell surface are shown. **(B)** Reconstruction and tracking of autophagosome progression from the pucta highlighted in (A). The AtEH1-GFP puncta appear associated with ATG8-labelled structures during early phagophore formation (bright puncta from 14s to 38s) and they stay associated during subsequent phagophore expansion. Scale bars: 5 µm (A), 1 µm (B).

**Fig. S3.**
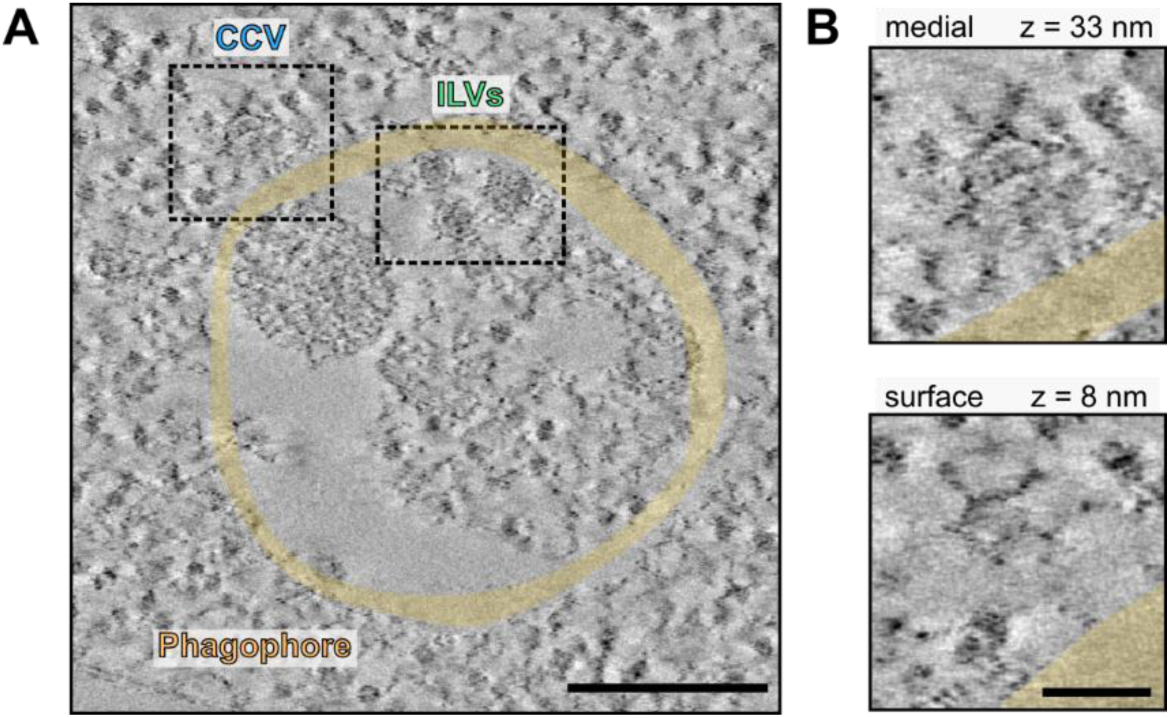
(Related to Fig. 2). CLEM-ET of a large amphisome. **(A)** Electron tomography reconstructed slices from a large autophagosome-MVB hybrid structure (amphisome) identified via CLEM from sorbitol + LatB treated *A. thaliana* root cells. (**B**) Close-up of a clathrin-coated vesicle attached to the autophagosome membrane. Distinctive clathrin triskelia pattern is visible on the surface view. Scale bars: 200 nm (a), 50 nm (b).

**Fig. S4.**
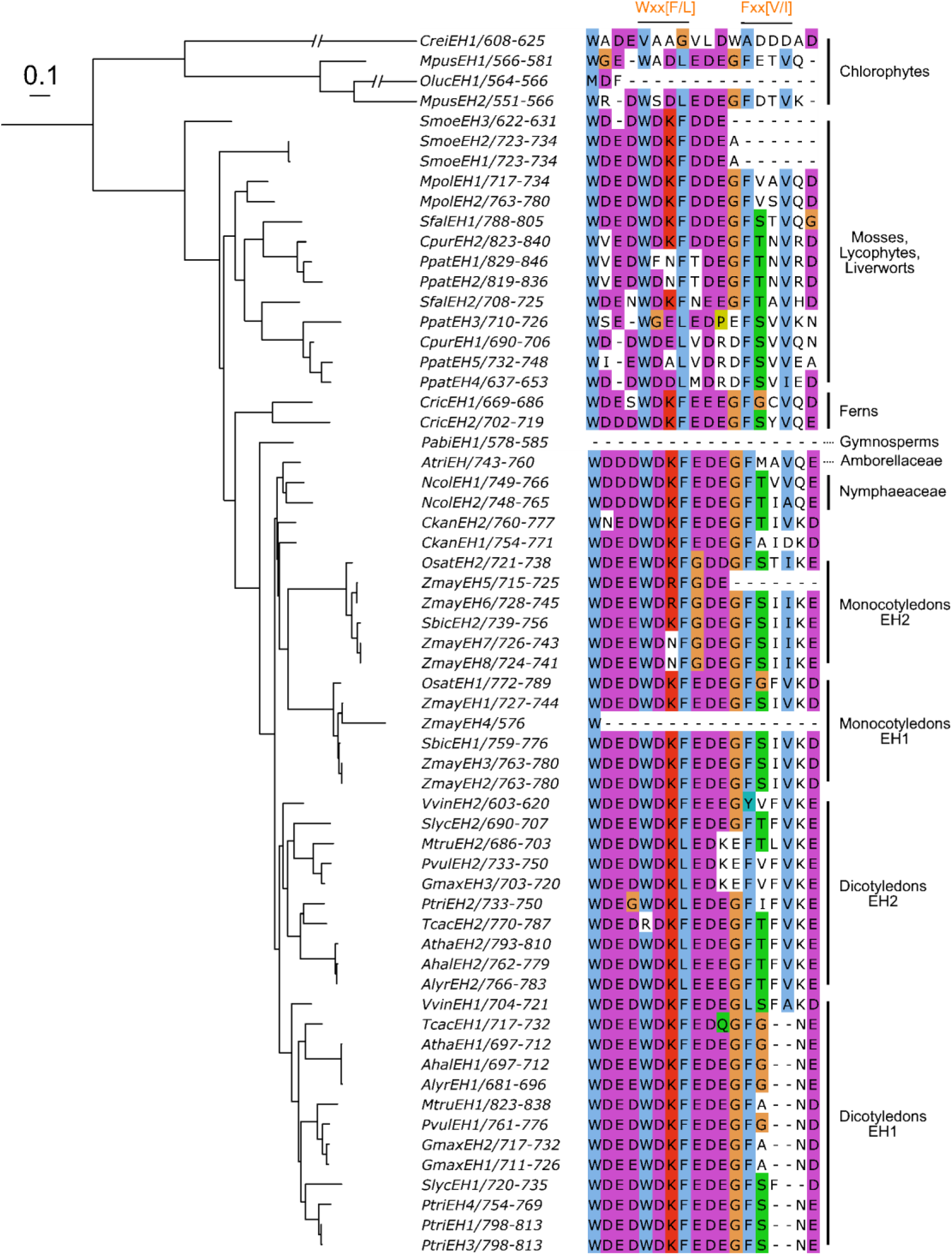
(Related to Fig. 4). Conservation of AIMs in plant EH/Pan1 proteins. Phylogenetic tree representing the maximum likelihood phylogeny of selected EH proteins. The phylogenetic tree was arbitrarily rooted to reflect phylogenetic relationships between chlorophyte and streptophyte lineages. The scale bar represents 0.1 amino acid substitution per site. The aligned region is in the IDR3 showing two AIMs: Wxx[F/L], present in most lineages, and Fxx[V/I], present in most lineages except for dicot EH1 proteins.

**Fig. S5.**
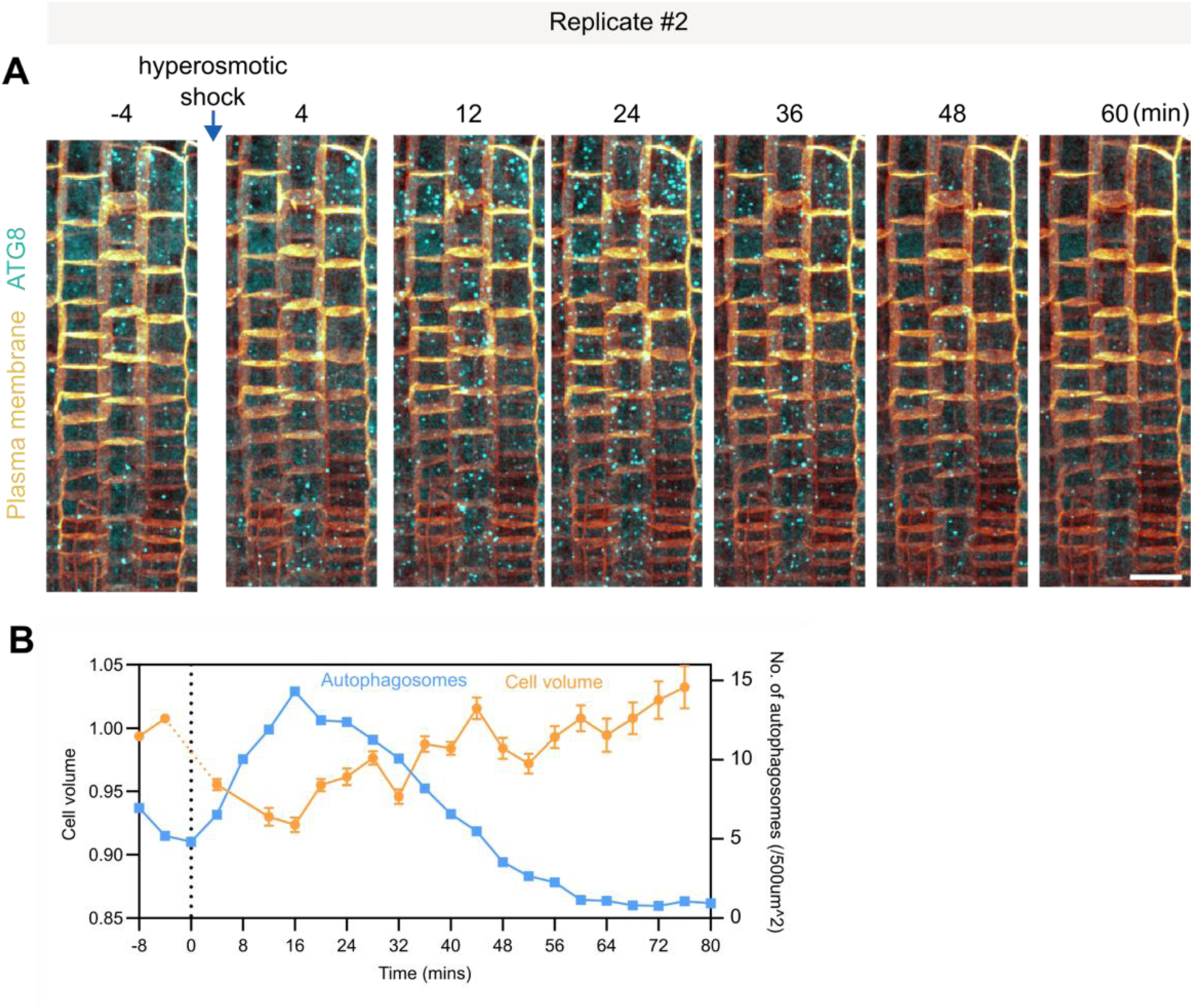
Autophagy and cell volume changes are interconnected following osmotic shock recovery. **(A-B)** Additional replicate of an osmotic shock experiment in *A. thaliana* roots. **(A)** Confocal images show *A. thaliana* root cells with autophagosomes (GFP-ATG8a), and plasma membrane (mCherry-SYP122) labelled before and after hyperosmotic shock. **(B)** Quantification of autophagosome puncta and cell volume. An inverse correlation is observed between autophagy induction and cell volume. Cell volume data is mean ± SEM; n = 32 cells. Scale bars: 20 µm.

**Fig. S6.**
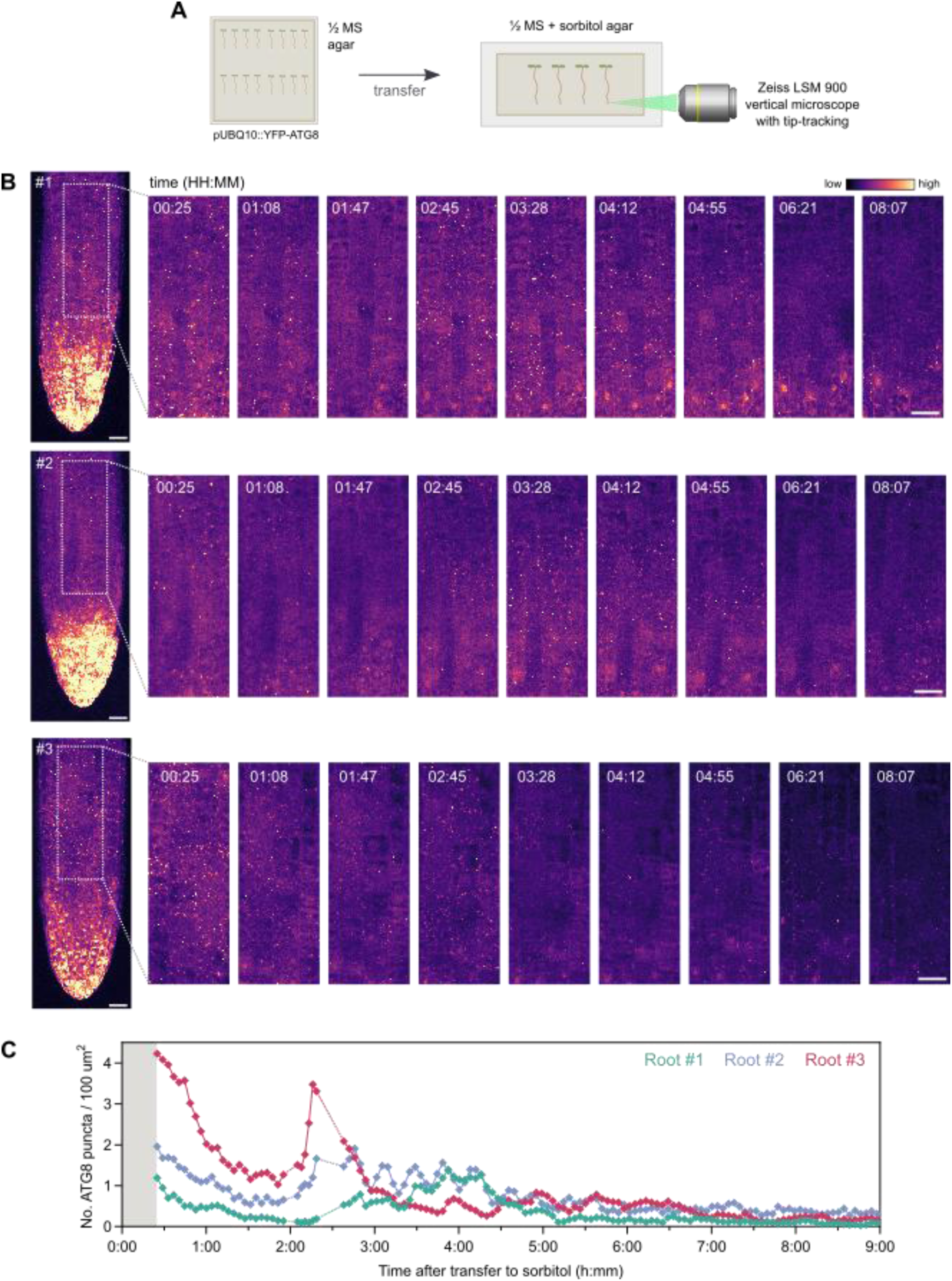
(Related to Fig 5): Autophagy dynamics during prolonged osmotic stress. **(A)** Schematic of the imaging setup. *A. thaliana* seedlings (pUBQ10::YFP-ATG8) were grown on ½ MS agar, transferred to chambered cover glass slides containing ½ MS + sorbitol, then imaged on a vertical microscope. Multiple root tips were imaged simultaneously every 4 minutes for 9 hours. **(B)** Maximum intensity projection of YFP-ATG8a signal in roots tips from three separate roots, and corresponding time-lapse images. **(C)** Quantification of ATG8a puncta over time showing that autophagy persists during prolonged osmotic stress beyond the initial osmotic shock response. Dashed lines indicate missing data points. Scale bars: 20 µm.

## Additional files

### Supplemental Video 1

Time lapse video showing autophagosomes (cyan, YFP-ATG8a), and plasma membrane (orange, mCherry-SYP122) before and after sorbitol induced hyperosmotic shock.

### Supplemental Video 2

Time lapse video showing autophagosome (YFP-ATG8a) induction during prolonged osmotic stress from seedlings transferred to agar containing ½ MS with 200 mM sorbitol.

**Table S1**. Mass spectrometry data

